# Distinct genetic pathways to music enjoyment

**DOI:** 10.1101/2024.04.04.588094

**Authors:** Giacomo Bignardi, Laura W. Wesseldijk, Ernest Mas-Herrero, Robert. J. Zatorre, Fredrik Ullén, Simon E. Fisher, Miriam A. Mosing

**Author notes:** Corresponding author: Giacomo Bignardi. These authors share last authorship (Fredrik Ullén, Simon E. Fisher, Miriam A. Mosing).

## Abstract

Humans engage with music for various reasons that range from emotional regulation and relaxation to social bonding. While there are large inter-individual differences in how much humans enjoy music, little is known about the origins of those differences. Here, we disentangled the genetic factors underlying such variation. We collected behavioural data on several facets of music reward sensitivity, music perceptual ability, and general reward sensitivity from a large sample of Swedish twins (*N* = 9,169). We found that genetic factors substantially explain variance in music reward sensitivity above and beyond genetic influences shared with music perception and general reward sensitivity. Furthermore, multivariate analyses showed that genetic influences on the different facets of music reward sensitivity are partly distinct, uncovering distinct pathways to music enjoyment and different patterns of genetic associations with objectively assessed music perceptual abilities. These results paint a complex picture in which partially distinct sources of genetic variation contribute to different aspects of musical enjoyment and open up new possibilities for using inter-individual differences to gain insights into the biology of a key aspect of human behaviour.

## Introduction

Music can evoke intense pleasure and induce various emotions ^1–4^, leading individuals from different cultures ^5^to actively seek out and engage with it. This human attraction to music has always been considered somewhat baffling ^6^and mysterious ^7^, leading many to ask why music has such power over humans ^8,9^. Oliver Sacks highlighted this conundrum in the opening of his beautifully written commentary, *The Power of Music*: “What an odd thing it is”, he wrote “, to see an entire species—billions of people—playing with listening to meaningless tonal patterns, occupied and preoccupied for much of their time by what they call ‘music’” ^9^. Despite the widespread power of music, however, it should also be noted that many people do not occupy themselves with music. Within human populations, there is indeed ample evidence that music-related cognition, from perceptual to affective-related processes, varies from one person to another ^10–13^.

Over the last decade, several studies have explored such differences between individuals in music-related traits and states to better understand the basis of human musicality ^14^. These studies show that differences in the ways individuals perceive, produce, or enjoy music correlate with neurobiological differences ^15–17^. For example, the study of individuals with lifelong musical pitch deficits underscores the relevance of brain connectivity patterns in distributed neural networks for conscious perception of music ^17^. Similarly, studies of differences in musical enjoyment highlight how interactions between cortical and subcortical brain regions support perceptual and affective processes that are fundamental for the experience of musical pleasure ^15,16,18–20^. Moreover, recent studies have started to uncover the roles of genetic factors in perceptual-motor processing of music ^21^ (e.g., the ability to synchronise with an external beat or recognise a melody) as well as in music production, such as levels of musical achievement ^22,23^. In general, these studies highlight complex gene-environment interplay ^24,25^ and the involvement of many DNA variants ^21^, each with a small effect (see ^26^).

Despite the many studies that have examined differences in music-related traits, still little is known about the genetic sources of differences in affective aspects of music processing and, in particular, the ability to enjoy music ^27,28^. A better understanding of such genetic effects will allow us to highlight how the ability to enjoy music is passed from one generation to the other and clarify the mechanisms linking genotypes, brains, and affect, providing a needed complementary perspective to resolve the conundrum of how “meaningless tonal patterns” can have such powerful effects on humans.

Here, we study individual differences in musical enjoyment, focusing on music reward sensitivity, a phenotype capturing how much individuals derive pleasure from music, as measured by the Barcelona Music Reward Questionnaire (BMRQ) ^12,16^. We used the BMRQ as it is a validated and reliable (e.g., one-year test-retest reliability, *R*_XX_(25) = .94, see ^12^) instrument that provides a fine-grained characterisation of individual differences in emotion evocation, mood regulation, music seeking, sensory-motor, and social reward facets of music enjoyment ^11^. Furthermore, it is a well-established psychometric tool in the music science literature, showing robust associations with affective experiences ^29–31^, cognition ^32–34^, physiology ^12^, and neurobiology ^15,16,35,36^. More specifically, we addressed the following three research questions:

1. To what extent are differences in music reward sensitivity explained by genetic variation?
2. To what extent do genetic effects influence music reward sensitivity above and beyond genetic effects shared with music perceptual ability and general reward sensitivity?
3. To what extent are genetic effects shared between the different facets of music reward sensitivity?

To address these questions, we utilised a large sample of deeply phenotyped monozygotic (MZ) and dizygotic (DZ) twins with available musicality data. We addressed the first question by estimating the heritability of music reward sensitivity using the classical twin design. We addressed the second question by applying multivariate twin modelling to estimate the genetic overlap between music reward sensitivity (BMRQ), music perceptual abilities based on a composite score of the melody, pitch, and rhythm scales of the Swedish Musical Discrimination Test (SMDT) ^13^, and general reward sensitivity, measured with the Behavioral Approach System Reward Responsiveness (BAS-RR) sub-scale ^37^, which has previously been shown to correlate with the BMRQ ^11,12,38^. The third question was assessed by testing if genetic effects are shared across facets of music reward sensitivity, consistent with a common genetic factor of music enjoyment, or whether, alternatively, genetic influences are distinct for each facet. Finally, we further extended the multivariate analyses at the facet level to explore associations with music perceptual abilities and general reward sensitivity.

## Results

### Sample and BMRQ descriptives

We utilised self-reported BMRQ data in a sample of 9,169 monozygotic (MZ) and same-sex and opposite-sex dizygotic (DZ) Swedish twins, with a mean (*M*) age of 51 years (standard deviation (σ) = 8 years, range from 37 to 64 years; see Table 1 for sample size split by sex and zygosity; see Methods for details on the cohort and zygosity identification). BMRQ total scores ranged from 20 to 100, with *M* = 71.20 and σ = 13.95. In line with previous studies, the BMRQ distribution was negatively skewed (skew = - 0.58; i.e., long tail of individuals with lower BMRQ total scores; see Supplementary Fig. 1). A confirmatory factor model showed acceptable fit for a model with a single latent music reward sensitivity factor capturing correlations between the five facets (CFI = .96, SRMR = .035).

**Table 1.**
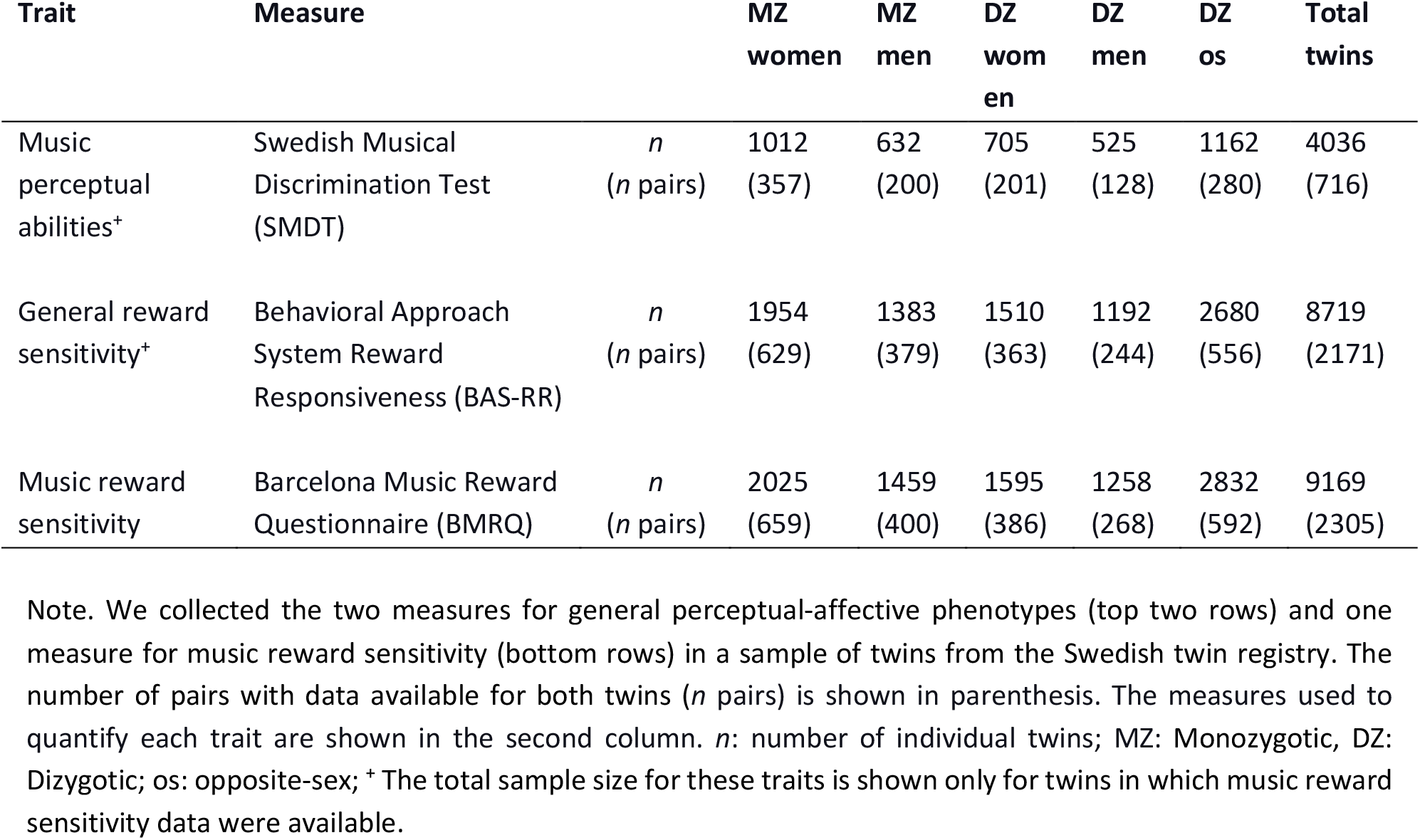
Numbers of monozygotic (MZ) and dizygotic (DZ) twin pairs for each trait.

**Fig. 1.**
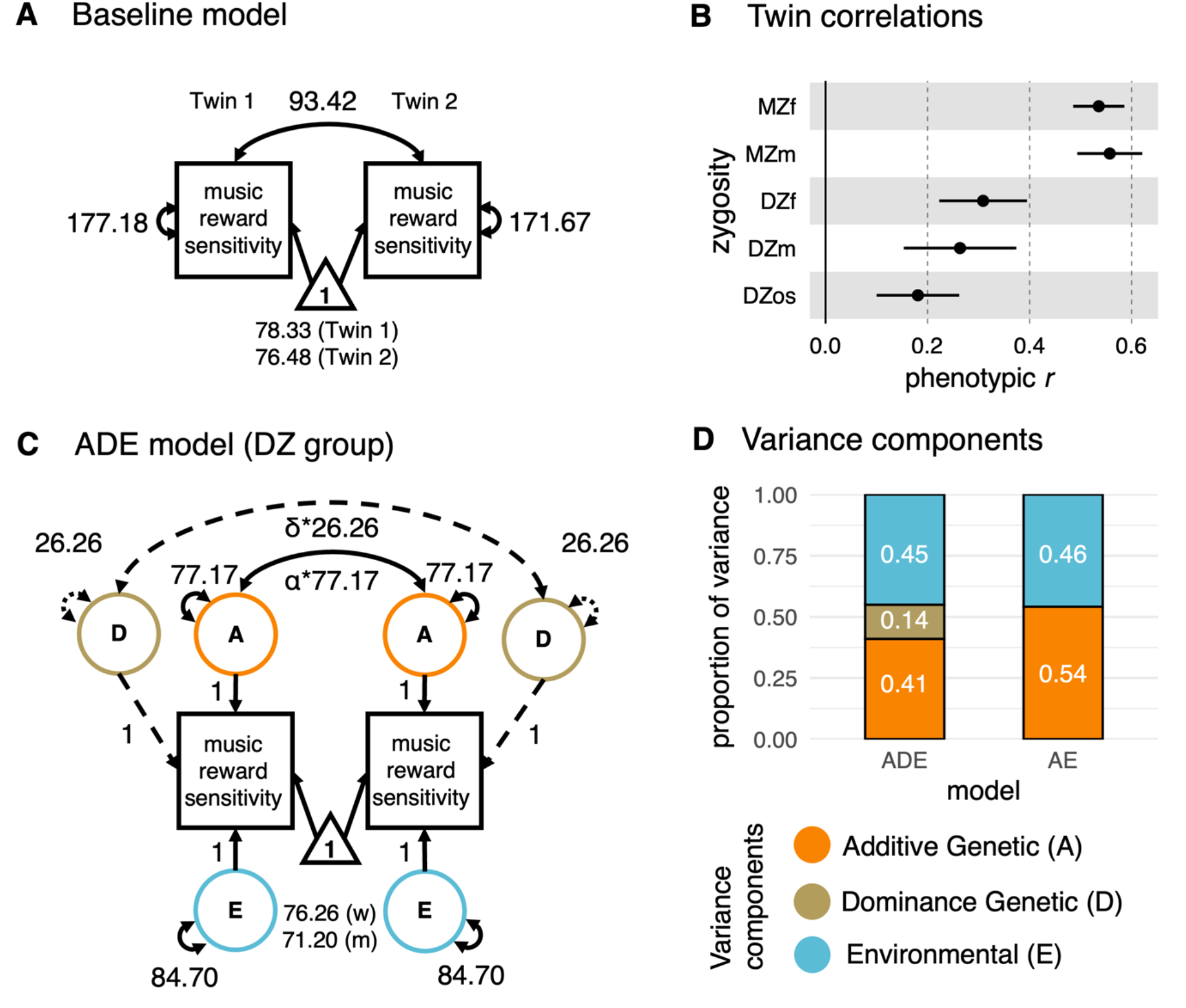
Music reward sensitivity is substantially heritable. (**A**) Baseline SEM to test for assumptions and further compare CTD-informed models fit; for simplicity only one group (MZ women) is shown. (**B**) Twin pair correlations grouped by zygosity and sex (women, w; men, m) extracted from the saturated model; note that MZ twin pairs are more than twice as similar in their music reward sensitivity as DZ twin pairs. The error bars represent 95% confidence intervals (CI). (**C**) The ADE model; note that we identified only A and E components as significant contributors to music reward sensitivity variability. α is the expected additive genetic relationship, and δ is the expected dominant genetic relationship between pairs (i.e., α = 1 or .5 δ =1 or .25, for MZ and DZ, respectively). (**D**) Estimated variance components from the final AE model indicated substantial heritability for music reward sensitivity. The left bar plot shows the estimates obtained from the full ADE model. *Notes on structural equation models*: *For simplicity, age is not included in the graphical representation of the model but is included as a covariate; Squares represent the measured phenotypes; Circles are the latent component; Double-headed arrows within circles, the variances associated with the latent components; double-headed arrows between circles covariances; the triangle, the phenotypic mean grouped by twin order (baseline model) and sex (ADE model) already adjusted for age; dashed elements, the component dropped after model comparison*.

### Genetic factors play a substantial role in music reward sensitivity

To estimate to what extent genetic effects (A: additive; D: dominance), the family environment shared between members of a family (C: common environment), and residual experiences unique to each individual (E: non-shared environment, including measurement error) influence music reward sensitivity, we use Structural Equation Modeling (SEM), informed by the Classical Twin Design (CTD). First, as a baseline for further model comparisons, we fit a univariate model to individuals’ BMRQ total scores (Fig. 1A; age and sex were accounted for). Assumptions of equality of means and variance across zygosities, twins within a pair, and sex were met (see Supplementary Table 1), except for the equality of means across sex: Consistent with previous literature ^39^, BMRQ scores were higher in women (*M* = 76.26) than in men (*M* = 71.20) (sex-constrained σ = 13.72; χ^2^(30)_Δ*df*_ = 300.54, *p* < 0.001). We, therefore, did not constrain means in subsequent models. Also consistent with previous results ^11,40^, age was negatively associated with overall BMRQ scores, although the effect was small, β_age_ = -0.03, (95% CI [- .05, -.01]), p = 0.004. Since the skewness of BMRQ scores was below 2 (see ^41^), all SEM analyses used the full-information maximum likelihood estimator. Analyses using alternative estimators, robust to departures from multivariate normality, did not change the findings; the results of these analyses are provided in Supplementary Note 1.

By comparing within-pair MZ and DZ correlations of BMRQ scores, we estimated the narrow-sense her itability (*h*^2^_twin_) of music reward sensitivity, i.e. the proportion of phenotypic variance in this trait which is explained by genetic variation ^42^. Twin correlations for music reward sensitivity were higher for MZ (*r*_MZ_ = .55, 95% CI [.51, .59]) than DZ (*r*_DZ_ = .24, 95% CI [.19, .29]) twins (Fig. 1B, see Supplementary Fig. 2). As the *r*_MZ_ was more than twice the *r*_DZ_, a model with additive and dominance genetics components (ADE) was fit (Fig. 1C). The ADE model reasonably fitted the data, as indicated by comparison against the baseline model (*χ*^*2*^(33) = 41.13, *p* =.16). However, a more parsimonious AE model, from which the D component was dropped, showed a better fit to the data (*χ*^*2*^(1)_Δ*df*_ = 1.63, *p* = .20). Therefore, the AE model was deemed the best fit for the data. The heritability for the BMRQ total score was substantial: *h*^2^_twin_ = .54 (95% CI [.51, .58]; Fig. 1D; see Supplementary Table 2 for details).

**Fig. 2.**
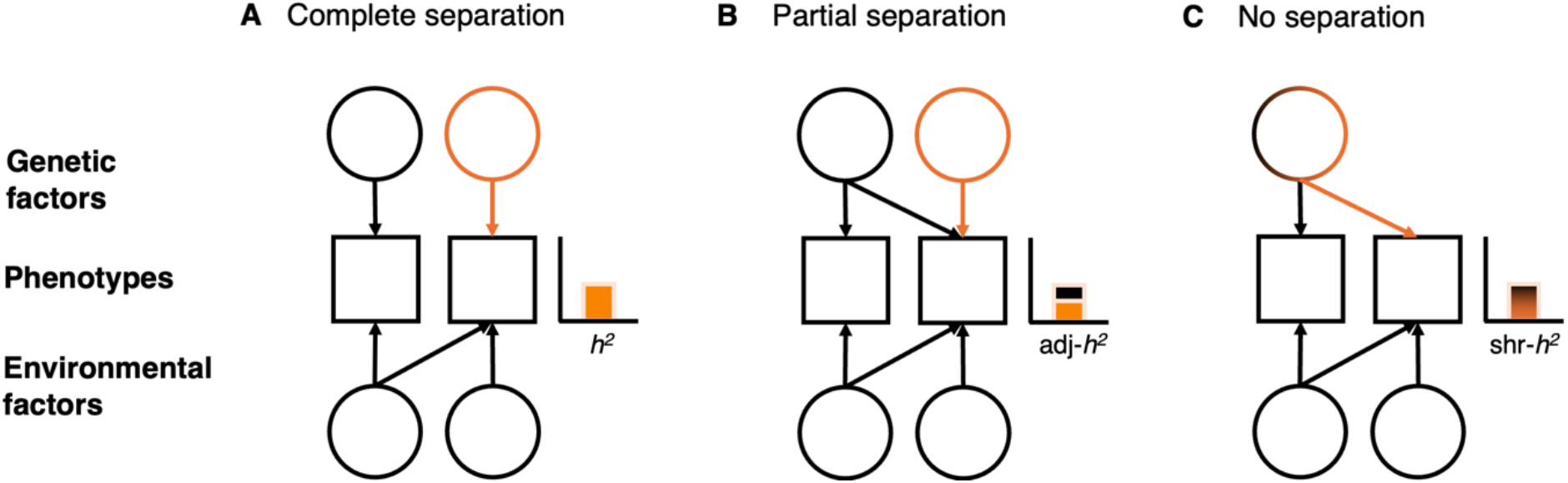
Schematic illustration of the sequential decomposition approach. The sequential decomposition of phenotypic associations employed to study unique and shared genetic influences. (**A**) The heritability (*h*^2^) of the second phenotype (orange bar) is fully separate from the genetic effect shared with the first. (**B**) In this case, after controlling for the *h*^2^ explained by the genetic effect shared with the first phenotype (black bar), an adjusted estimate (adj*-h*^2^, remaining orange bar) is still substantial. (**C**) Here, *h*^2^ is completely shared (shr-*h*^2^) between the two phenotypes. For simplicity, only two traits are shown.

### Music reward sensitivity is influenced by genetic factors above and beyond genetic influences shared with music perceptual abilities and general reward sensitivity

To better understand the nature of genetic effects contributing to music reward sensitivity, we tested whether the genetic influences on BMRQ were partly shared with other related traits, such as music perceptual abilities and general reward sensitivity. For this purpose, we used a multivariate sequential decomposition approach, which allowed us to discriminate between three possible outcomes, as illustrated in Figure 2. Genetic effects on music reward sensitivity could be either fully (Fig. 2A) or partly (Fig. 2B), separate from genetic effects on music perceptual abilities or general reward sensitivity. Alternatively, they could be fully shared (Fig. 2C) and hence entirely accounted for.

First, we revealed and confirmed that there were significant phenotypic correlations between music reward sensitivity and music perceptual abilities and general reward sensitivity ^11,12,38^, respectively (*p* <.001; Fig. 3A; correlations were estimated from a sample of only one twin per pair, to avoid sample dependence; estimates were similar in the other twins, see Supplementary Fig. 3 for details). To simultaneously accommodate the three phenotypes, we employed a tri-variate sequential decomposition. This analysis indicated partial separation of genetic (and environmental) factors influencing the three variables (Fig. 3B; see Supplementary Table 3 for coefficient estimates). The *h*^2^_twin_ of music reward sensitivity adjusted for music perceptual abilities and general reward sensitivity was adj-*h*^*2*^_*twin*_ = .38 (95% CI [.33,.43], Fig. 3C). Thus, of the total variance in music reward sensitivity explained by genetic factors (*h*^*2*^_*twin*_ = .54), around 70% (95% CI 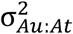 = [.63,.78]) was unique to this trait. Only the remaining 30% was shared with genetic effects on music perceptual abilities and general reward sensitivity, explaining 12% and 18% of the total genetic variance in music reward sensitivity, respectively. Environmental influences shared across phenotypes, which reached significance only for general reward sensitivity (*p* <.001), explained only 2% of the total variance in music reward sensitivity (see Supplementary Note 2).

**Fig. 3.**
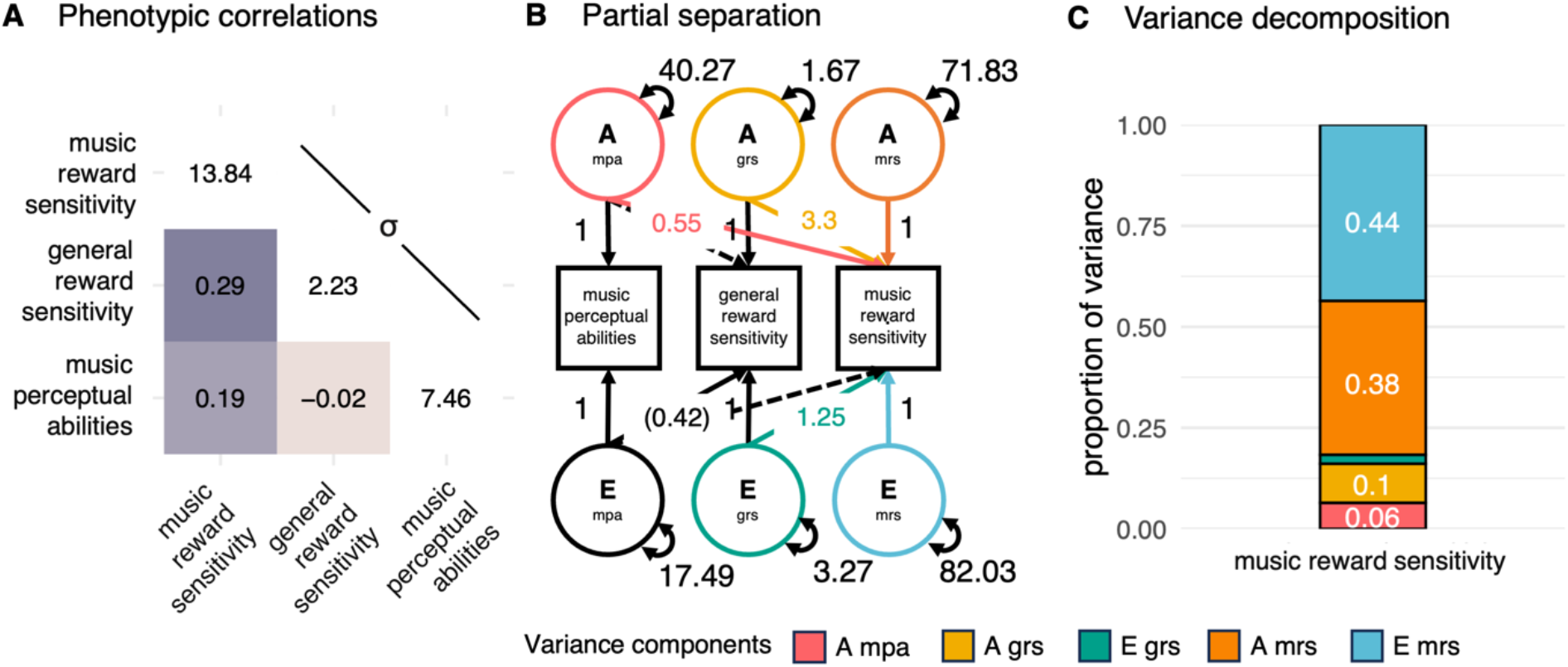
Genetic effects on music reward sensitivity are partly separate from music perceptual abilities and general reward sensitivity. (**A**) All cross-phenotypic correlations with music reward sensitivity were significant (all *p*<.001). On the diagonal, the standard deviations (σ). (**B**) Sequential decomposition of the significant contributions to music reward sensitivity; note that the environmental path from music perceptual abilities to music reward sensitivity is not significant, indicating only common genetic causes. Between parentheses, the significant path from the E component to general reward sensitivity (*p* = .03) (**C**) Variance decomposition shows that genetic factors explain individual differences in music reward sensitivity (in orange) well beyond shared genetic factors associated with known general perceptual and affective processes (in red and yellow, respectively). The variance components here indicate the proportion of variance explained by the respective components. mpa: music perceptual abilities; grs: general reward sensitivity; mrs: music reward sensitivity. *Notes on structural equation models*: *one-headed arrow represents regression paths partitioned in additive genetics and unique environmental paths; dashed one-headed arrows represent non-significant paths. Other abbreviations and symbols are as in Fig. 1*.

### Genetic pathways to the different facets of music-reward sensitivity are partly distinct

Having shown that music reward sensitivity has substantial heritability and is partly genetically separate from relevant general perceptual-affective processes, we went on to test whether the pattern of genetic correlations across facets is consistent with an overarching one-genetic-factor solution for music reward sensitivity (Fig. 4 A-C). If largely distinct genetic pathways influence the different facets of music enjoyment, a one-genetic-factor solution would not be supported. This scenario can be modelled as a multivariate correlated factor solution, which solely allows for genetic and environmental pairwise correlations (Fig. 4B). If, on the other hand, there is a common genetic source of different aspects of musical enjoyment, we would expect underlying genetic sources of variability to be mostly shared across different facets (Fig. 4C, see ^43^). This latter scenario can be instead modelled as a multivariate hybrid independent pathway model (see ^44^). Here, along with distinct genetic effects over single facets, an extra additive genetic common factor is modelled to capture shared genetic effects across all facets. For ease of interpretation, we will hereafter refer to the model depicted in Fig. 4B as the distinct factor solution and the model depicted in Fig. 4C as the common-genetic factor solution.

**Fig. 4.**
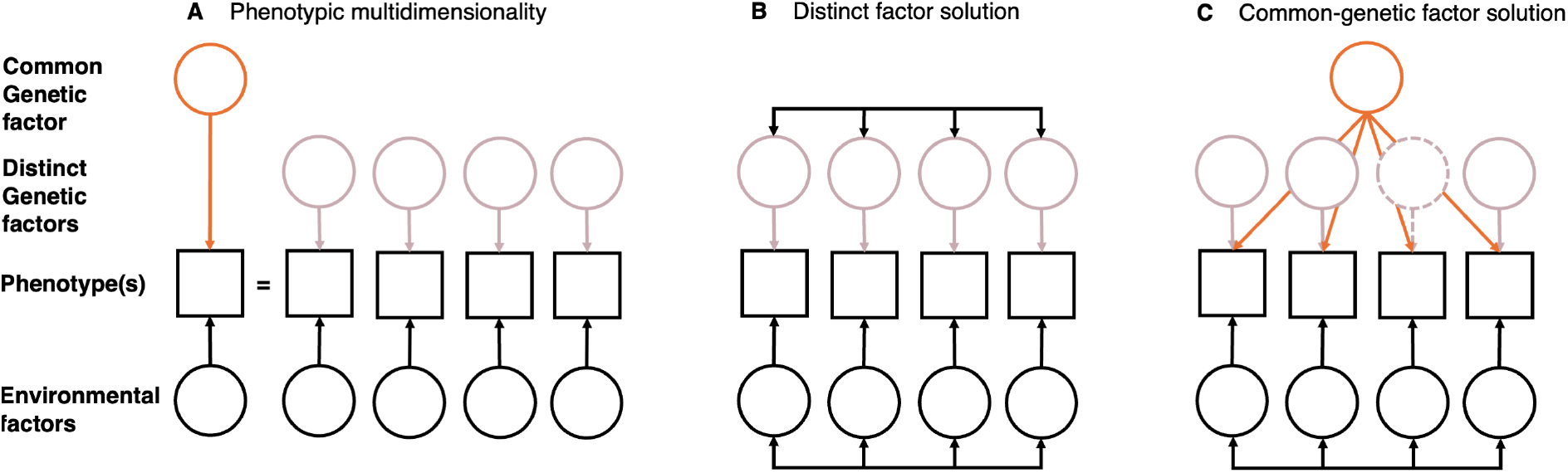
Schematic illustration of multivariate models employed to quantify distinct and common genetic factors. (**A**) The phenotype is decomposed into its constituent facets. (**B**) The first solution includes distinct genetic factors with a simple description of all possible genetic and environmental covariances. (**C**) A common-genetic factor solution is applied by assuming a genetic factor that captures the genetic covariances across facets. The common latent genetic factor (in orange) could explain all the genetic variance associated with one facet (e.g., dashed circle). (Double-headed arrows are compressed to avoid cluttering.) Figure inspired by ^45^.

Since the common-genetic factor solution is a constrained version of the distinct factor solution, model comparisons can be used to test whether a common-genetic factor of music reward sensitivity facets shows a better fit to the data. While both models fit the data well (CFI= .988, SRMR = .048, and CFI = .981, SRMR = .061, respectively; See Supplementary Table 4), the common-genetic factor worsened the fit of the distinct factor solution (χ^2^(5)_Δdf_ = 129.61, p < 0.001;). This implies that the distinct factor solution is a more appropriate description of the structure of the genetic effects compared to the common-genetic factor solution. (Fig. 5A-B; See Supplementary Note 3 for more details).

**Fig. 5.**
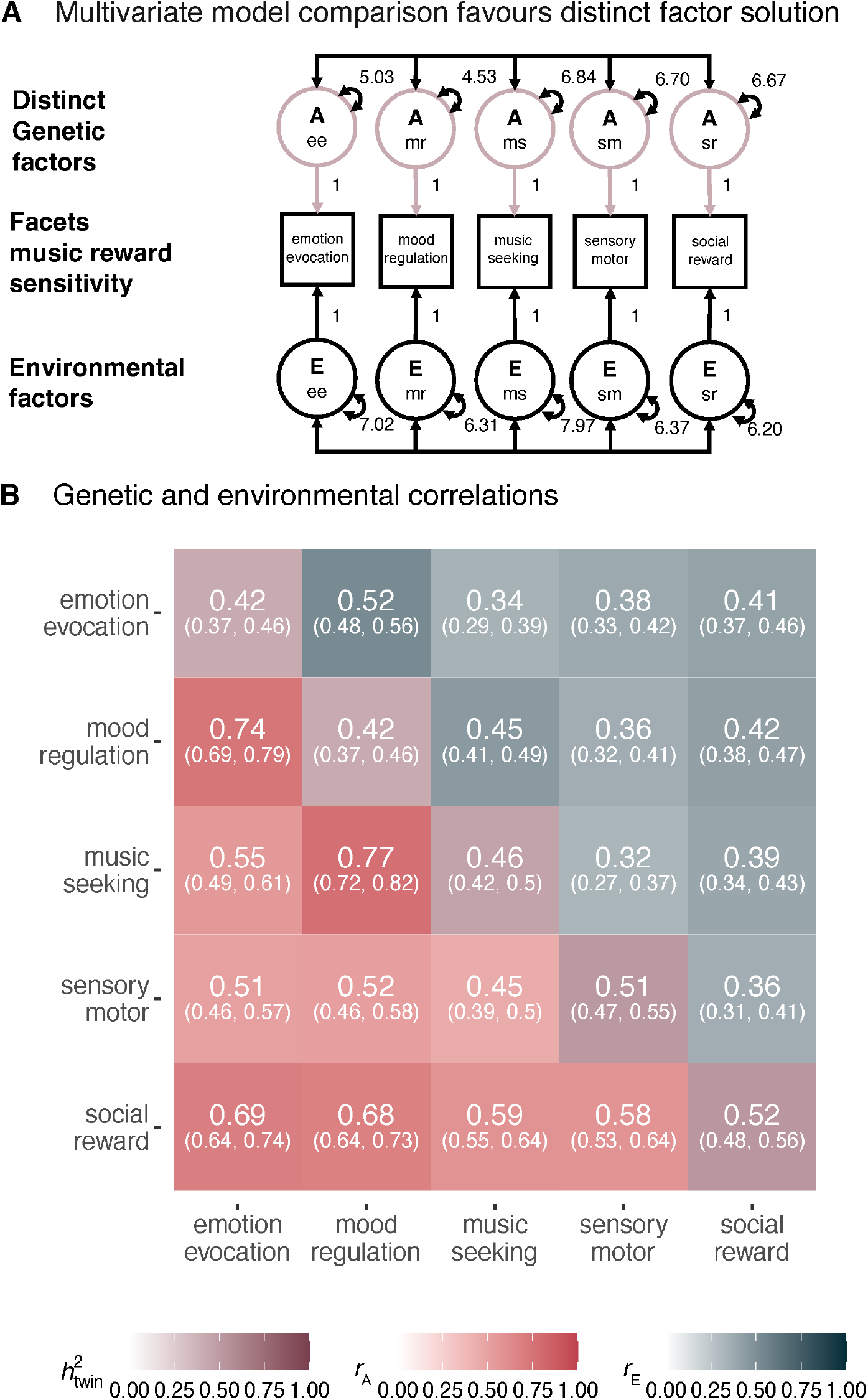
Genetic heterogeneity between distinct musical affective phenotypes. (**A**) Simplified distinct factor solution of music reward sensitivity facets. (**B**) Genetic effects on music reward sensitivity are partially heterogeneous. Matrix extracted from the correlated factor model. Additive genetic (*r*_A_) and environmental correlations (*r*_E_) are shown below (red) and above (blue) the diagonal, respectively; numbers on the diagonal show heritability estimates. Numbers in parentheses are 95% confidence intervals. Note that genetic correlations are far from 1, suggesting that music reward sensitivity has multiple genetic sources. Phenotypic correlations can be found in Supplementary Fig. 4. *Notes on structural equation models*: *double-headed arrows between circles represent A and E covariance between facets. Other abbreviations and symbols are as in Fig. 1 and 3*.

### Exploratory analyses reveal that social reward shares substantially more genetic variance with music perceptual abilities than the other facets

Having shown that genetic influences are partially distinct between music-reward sensitivity facets, we further explored such genetic heterogeneity by fitting two additional multivariate distinct factor solutions to data on music reward sensitivity facets, with music perceptual abilities and general reward sensitivity added to the models (Fig. 6A). Additive genetic correlations (*r*_*A*_) between music reward sensitivity facets and music perceptual abilities varied widely (range *r*_*A*_ = .15 to *r*_*A*_ =.49; Fig. 6B), with differences between the *r*_*A*_ values (Δ*r*_*A*_) being significant (Supplementary Table 5). Specifically, the Δ*r*_*A*_ estimates were significantly higher for the social-reward facet of music reward (*r*_A_ = .49, 95% CI [.42; 56]) than for any other facet (range Δ*r*_*A*_ from .19 to .39, all *p* < .001). In comparison, *r*_*A*_ obtained from the model fit to general reward sensitivity data were similar across facets (range *r*_*A*_ = .29 to *r*_*A*_ =.36; Fig. 6C) and did not significantly differ (all *p* > .05). These observations further strengthen the evidence that different aspects of music reward show functionally relevant genetic heterogeneity.

**Fig. 6.**
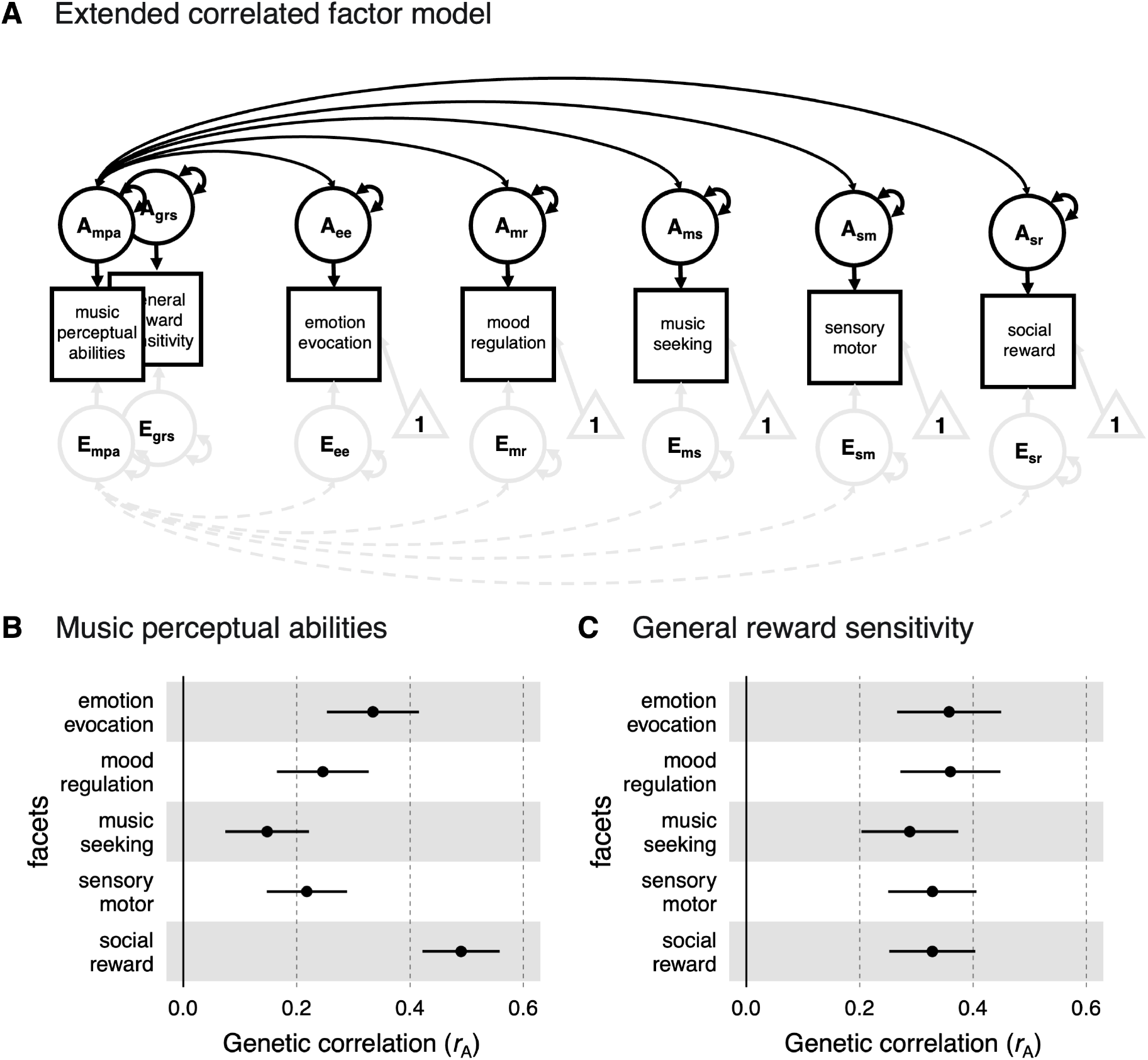
Genetic heterogeneity between distinct musical affective phenotypes. (**A**) The correlated factor model extended to estimate genetic correlations with music perceptual abilities and general reward sensitivity. (**B-C**) Magnitude of the genetic correlations (*r*_*A*_) between facets of music reward sensitivity and music perceptual abilities (B) and general reward sensitivity (C). Error bar represents 95% CI. *Notes on structural equation models*: *For simplicity, all pairwise covariances are not included but are present in the model—other abbreviations and symbols are as in previous Figures*.

## Discussion

Our understanding of why “meaningless tonal patterns” ^9^ have such powerful effects on humans can benefit tremendously from the study of inter-individual differences. Here, by exploiting a large and deeply phenotyped Swedish twin sample, we found that music reward sensitivity has substantial heritability. Most of this genetic variance influences music reward sensitivity independently of music perceptual abilities and general reward sensitivity, suggesting genetic variations influence music reward sensitivity not only via other general perceptual-affective processes. Furthermore, our findings reveal considerable genetic heterogeneity behind different facets of music reward sensitivity. Although all facets show heritability estimates of a similar magnitude (between 42% and 52%) and are genetically correlated (between .45 and .77), the results do not support a single (genetic) dimension of musical enjoyment. Instead, these findings are consistent with musical enjoyment being built upon genetically interconnected yet partly distinct parts. Extended multivariate analyses further strengthened these results by showing that music perception shows stronger genetic correlations with social bonding than other facets of music reward, indicating functionally relevant genetic heterogeneity.

Despite answering a long-standing question ^27,28^, the finding that music reward sensitivity is to some extent heritable is not surprising in light of the fact that virtually every human trait is at least partly genetically influenced ^46,47^. Yet, the finding of notably high heritability for music reward sensitivity gives hope for molecular genetic studies to answer questions about genetic underpinnings of musicality in general and musical affect in particular. Prior studies of individual differences in music reward sensitivity ^12,15,16,48^ have had far-reaching implications for our knowledge of biological pathways implicated in perceptual-affective processes ^18–20^. These studies have shown that individual differences in music reward sensitivity are associated with variation in functional and structural connections between two systems. The first includes the auditory cortex and its pathways involved in perceptual analysis, feature encoding, and working memory. The second, the reward system, encompasses the striatum, orbitofrontal cortex, and ventral tegmental area and is involved in pleasure, salience, and learning ^1,2,16,19,20,49^. These neurobiological mechanisms could provide a potential substrate for the genetic influences identified in the present study. Therefore, an important question for future studies is to investigate whether variability in structural and functional properties of the relevant brain networks, and their interactions, may mediate the genetic effects on the ability to enjoy music, thus furthering our overall mechanistic understanding of a key aspect of human affect.

A complementary genetic perspective on music reward sensitivity could be a particularly fruitful strategy to better understand human musicality and affect because we found genetic influences to be primarily separate from other relevant perceptual-affective processes, such as music perceptual abilities and general reward sensitivity. The dissociation between the genetics of music reward sensitivity and general perceptual and reward processing mirrors the finding that specific musical anhedonia, i.e., blunted or absent hedonic responses from music stimuli, exists in the absence of any perceptual or generalised reward deficit ^12,16^, yet contrasts other findings suggesting sensitivity to intrinsic rewards to be domain general ^50^. This implies that genetic variance associated with music reward sensitivity, beyond perceptual and general reward processing, can be used to better disentangle and understand the mechanisms involved in sensory-specific experiences of enjoyment.

The partial separation between genetic effects on perception and enjoyment also opens up the possibility that genes influencing music perception and enjoyment may have been a distinct target for natural selection during evolution ^51^. The finding implies that genetic variation between people may be used to dissect the evolutionary trajectories of different aspects of human musicality. Along these lines, a further question of interest becomes whether genetic variants, which are more specifically associated with music enjoyment, are also enriched in genomic regions of evolutionary interest ^52,53^.

Here, we did not find support for a single overarching genetic factor of music reward sensitivity. On the contrary, we found several distinct genetic pathways to music enjoyment. This result aligns with general views of musicality as “built upon a suite of interconnected capacities, of which none is primary” ^54^. Our results demonstrate that such heterogeneity is seen even when zeroing in on one hypothesised core feature of musicality: enjoyment. We show that music reward sensitivity is itself not a monolith and that different facets of this trait are influenced by partly different genetic pathways; these facets range from the ability to experience emotion and get chills to the rewarding aspects of social bonding through music. Our results thus may challenge the epistemological status of music reward sensitivity as a latent causal factor ^43,55,56^, as a latent factor is unlikely to hold unless a common-genetic factor solution holds (for additional conditions, see ^43^).

Our final exploratory analysis provides a direct example of the implications such a shift in perspective might have on the study of human behaviour and affect. When dissecting the genetic effects at the level of the facets of music reward sensitivity, novel insights emerge. Our findings indicate that music perceptual abilities are genetically more strongly correlated with rewards of social bonding through music. This could be seen as in line with the social bonding hypothesis, which states that “core biological components of human musicality evolved as mechanisms supporting social bonding” ^57^. This was not the case for the association between music reward and general reward sensitivity, which were relatively similar across different facets. Furthermore, shared additive genetic variation entirely explained the association between music perceptual abilities and social reward, suggesting shared biological components to be at play. These results highlight how acknowledging the genetic heterogeneity of music reward sensitivity might reveal associations that might have been otherwise unnoticed. (For a detailed discussion on consequences for other well-studied conditions, such as musical anhedonia ^12,15,16^, we refer to Supplementary Note 4.)

Notwithstanding such functionally relevant genetic heterogeneity, we also found genetic overlap between the facets, suggesting genetic effects over music reward sensitivity are also partially shared. This finding is important as some degree of genetic overlap across facets of music reward sensitivity is needed to better understand the biology of music enjoyment as a whole. Further studies could test whether these genetic effects underlie other auditory phenomena, such as pleasure derived by timbre in sounds, which has been shown to correlate homogenously across facets of music reward ^29^ or other broader aspects related to human affect, such as aesthetic sensitivity ^58^.

Finally, the absence of shared environmental effects on music reward sensitivity (at least under the assumption of the classical twin design, see below) aligns with many other complex traits, including those related to musicality ^26,46^. Yet, it contrasts with findings on some musicality traits, such as musical achievement ^23,24^ or singing abilities ^59^, for which modest effects of shared environment have been found using similar designs. The lack of shared environmental effects for some traits but not others suggests that different aspects of musicality, namely producing music and enjoying music, might follow different patterns of intergenerational transmission. The likely absence of shared environmental effects may imply only a small, if present, passive gene-environment correlation (e.g., genotypes associated with music reward sensitivity in the parents influence the children via the environment the parents provide and the genes they pass on to their children, see ^60^). This is crucial because passive gene-environment correlations would complicate future efforts to detect direct genetic effects on music reward sensitivity by, e.g. confounding direct genetic effects with indirect effects caused by the environment that the parents provide to their children (see ^60,61^ for a detailed discussion). Recent efforts to better understand the genetic architecture of complex traits focus on deconstructing indirect sources of heritability, which inflate estimates of genetic effects and confound the possible inferences that can be obtained from downstream analysis of genome-wide-derived estimates ^61–63^. Our findings suggest that music reward sensitivity, or rather its constituent facets, may be especially promising for facilitating discoveries of direct molecular genetic effects on music enjoyment.

As with every other twin-informed study ^42^, our work depends on a number of assumptions. In the Methods section, we highlight these assumptions and what violation of each entails. One critical assumption is the equal environment assumption, which states that environmentally caused differences between twins within a pair are the same across zygosities. An additional assumption is the lack of gene-by-shared environment interaction, which could lead to an underestimation of the variance of the C component. For example, additive genetic effects associated with music reward sensitivity might vary within different musically enriched environments. However, we also note that the equal environment assumption is not violated if different zygosities experience more similar or dissimilar environments due to genetic differences. On the contrary, this is to be expected if evocative and active gene-environment correlations are at play, which seems likely for traits related to music enjoyment. Such gene-environment correlations would not inflate *h*^*2*^ estimates. Still, they would change their interpretation as they could reflect, for example, a more complex causal chain that leads individuals to seek or be exposed to environmental changes that, in turn, influence the phenotype, resulting in processes such as niche picking ^64,65^.

At the same time, our study also exploits one of the fundamental strengths of the CTD —the possibility to estimate genetic effects on deep phenotypes, such as objectively assessed music perceptual abilities and the full BMRQ, which are notoriously difficult to obtain in large genetically informative samples ^66^. In light of the limitations and the strengths of the CTD, our *h*^*2*^_*twin*_ can be considered both an upper bound for the *h*^*2*^ (within an environment, a population, and at a given time) and provide valuable benchmarks for the total effect of DNA variation ^42,64,67^ of music reward sensitivity and facets, above and beyond perceptual-affective processes. These findings, as discussed in length above, generate novel insights and pave the way for future research on the genetics of music enjoyment and human affect.

## Conclusions

Musicality is the capacity that allows individuals of a species to perceive, generate, and enjoy music ^14,54^. Much has been said about the sources of the considerable inter-individual variation in music perception, production, participation, and achievement. Yet, relatively little has been written on the genetic contribution to what makes individuals differ in their capacity to enjoy music. Here, we add a new piece to the puzzle of why music has such powerful effects on humans. We show that genes influencing our ability to enjoy music are largely distinct from genes involved in other, more general aspects of perceptual and affective processing. Further, we reveal that genetic pathways to music enjoyment are partially distinct and that the genetic overlap between music perceptual abilities differs between different facets of music reward. In summary, the findings highlight the complex and multifaceted nature of music enjoyment and its genetic underpinnings, paving the way for further studies of the evolutionary origins and genetic and neural mechanisms for a key aspect of human affect.

## Methods

### Sample

#### Swedish Twin Registry: Screening Twin Adults Genes and Environment (STAGE)

Participants were twins recruited from the Swedish Twin Registry ^68^. Twin zygosity was determined by questionnaire data, which, when compared to genotypes, has been shown to be 99% accurate in the Swedish Twin Registry ^69^. The twins included in this study took part in two large recent waves of online data collection on music, art and cultural engagement. In 2011 and then again in 2022, a total of 32,000 adult twin individuals were invited from the STAGE cohort born between 1959 and 1985, of which around 11,500 participated in the first wave and then around 9,500 in the latest wave. More details on the survey can be found in Ullén et al. ^13^. Participants took the Swedish Musical Discrimination Test (see below) in the first wave and responded to the Behavioral Approach System and Barcelona Music Reward Questionnaire in the second wave of data collection. A full description of the twin sample across waves of data collection can be found in Table 1, including *n* of twins for which we had both data available, stratified by the zygosity and the sex of the twins; for both waves of data collection, informed consent was given by each participant before data gathering began. Both studies were approved by the Regional Ethical Review Board in Stockholm (Dnrs 2011/570-31/5, 2012/1107/32, 2021-02014, 2022-00109-02, 2020-02575).

### Primary measure

#### Barcelona Music Reward Questionnaire (BMRQ)

The Barcelona Music Reward Questionnaire (BMRQ) is a psychometric tool used to assess musical anhedonia ^12,16^ and, more generally, music reward sensitivity ^11^, which has previously been validated across many cultures ^11,70–72^. It comprises 20 self-report items, with five response options, ranging from completely disagree to completely agree. After recoding response items to numeric options (1 to 5), with two out of 20 items being reverse coded, we used the sum score of the BMRQ as a measure of music reward sensitivity (score range from 20 to 100). Following the original five-factor structure ^11^, we also created sum scores of the five known facets of music reward sensitivity ^28^: (1) Emotion-evocation - the degree to which individuals get emotional, experience chills, and even cry when listening to music; (2) Mood regulation - the degree to which individuals experience rewards from relaxing when listening to music; (3) Musical seeking – the pleasure associated with the discovery of novel music-related information; (4) Sensory motor – the rewards obtained from synchronising to an external beat or dancing; (5) Social reward – the rewards of social bonding through music. Additional details are given in Supplementary Note 5.

### Secondary measures

#### Behavioral Approach System Reward Responsiveness (BAS-RR)

The Behavioral Approach System (BAS) scale is included in the Behavioral Inhibition System (BIS)/BAS questionnaire, a validated psychometric tool to assess inter-individual differences in two general motivational systems ^37,73^. The BAS-Reward Responsiveness (BAS-RR) scale, in particular, assesses inter-individual differences in the ability to experience pleasure in the anticipation and presence of reward-related stimuli and predicts general psychological adaptive functioning ^74^. It comprises five items, with four response options for each. BAS-RR is obtained by the sum score of the five items after the numerical conversion of the responses (1-4). Additional details are given in Supplementary Note 6.

#### Swedish Musical Discrimination Test (SMDT)

The Swedish Musical Discrimination Test (SMDT) is a test that has good psychometric qualities for individual abilities in auditory perceptual discrimination of musical stimuli ^13^. It comprises three subtests: melody, rhythm, and pitch. A brief description of each test is given below (see ^13^ for more details).

#### Melody

This subtest used isochronous sequences of piano tones as stimuli. Tones ranged from C4 to A#5, played at 650 ms intervals (American standard pitch; 262–932 Hz). The number of tones increased from four to nine during the subtest progression. For each of the six stimulus lengths, there were three items. The two stimuli in an item were separated by 1.3 s of silence. The pitch of one tone in the melody was always different in the second stimulus. Participants had to identify which tone was different.

#### Rhythm

In this subtest, each item included two brief rhythmic sequences of 5-7 sine tones, lasting 60 ms each. The inter-onset intervals between tones in a sequence were 150, 300, 450, or 600 ms. The two sequences in an item were either identical or different, and separated by 1 s of silence. The participant had to determine whether the two sequences were the same or not.

#### Pitch

The pitch subtest used sine tones with a 590 ms duration as stimuli. In each item, two tones were presented, one of which always had a frequency of 500 Hz. The frequency of the other tone was set between 501 and 517 Hz. The order of the two tones varied randomly, with tones separated by a 1 s silence gap. Participants had to identify whether the first or the second tone had the highest pitch. The item difficulty was increased progressively by gradually making the pitch differences between the tones smaller.

### Analyses

#### Factor Analysis

To confirm the BMRQ‘s sum score as an appropriate measure of music reward sensitivity in the Swedish sample, we ran a one-factor Confirmatory Factor Analysis (CFA) on the five facets of the Swedish version of the BMRQ. CFA was run on one twin per pair, using the lavaan::cfa() function, to avoid sample dependence.

#### Classical twin design (CTD)

The CTD allows the estimation of additive (A) or dominance (D) genetics, shared environmental (C), and residual source (E) of phenotypic variance (σ_A_^2^, σ_D_^2,^ σ_C_^2^, and σ_E_^2^, respectively). This is possible given the expected phenotypic resemblance of monozygotic (MZ) and dizygotic (DZ) twins. MZ arise from the same fertilised egg and thus are ∼100% genetically similar (with minimal deviations from expected genetic similarity, see ^75^); DZ arise from separate egg cells and thus, as ordinary siblings, share on average 50% of their segregating genes. Furthermore, when both twins of a pair are raised in the same household, MZ and DZ share 100% of their common environment. Finally, by definition, remaining deviations from the expected values inferred by additive, dominant, and shared environmental effects represent unique environmental influences and measurement errors. Therefore, E is not shared between twins within a family. Under a set of assumptions, including no epistasis (gene-by-gene interaction, see ^76^), the covariance of MZ twin pairs is then equal to:

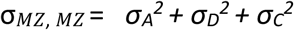

While the covariance of DZ twin pairs is equal to:

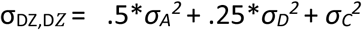

Given that the variance and covariance are measured between twins within families, it is possible to specify a multigroup structural equation model and estimate three out of four variance components. The decision of which parameters to include in the model (e.g., A, C, E, or A, D, E) is purely based on twin covariances, which are extracted from the baseline phenotypic model (for details on the baseline model, see below), and biological plausibility. If σ_*MZ, MZ*_ > 2^*^σ_D*Z*, D*Z*_, then D is expected to contribute to the phenotypic variance, and, therefore, an ADE model is specified (note that DE models are not biologically plausible). Otherwise, an ACE model is fit to the data.

#### CTD assumptions

The estimates from the CTD are unbiased under a set of assumptions. First, the CTD assumes equal environments between the twins. In other words, it assumes that similarities between twins caused by the environment are the same for both zygosities. Suppose, instead, MZ experiences their environment more similarly than DZ due to environmental, not genetic, causes. In that case, the estimate for the genetic variance will be upwardly biased (i.e., 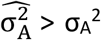) Note that the equal environment assumption is not violated if MZ experiences their environment more similarly than DZ due to genetic differences. The latter case would instead result in active gene-environment correlations that are still consistent with the estimate of the variance components. The second assumption is that the phenotypes of the parents of the twins’ are uncorrelated (i.e., random mating, also known as panmixia ^77^). If the covariance between two parental phenotypes, *p*_*1*_ and *p*_*2*_, is different from 0, *σ*_*P1,P2*_ ≠ 0, then the shared environmental variance might be upwardly biased (i.e., 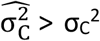) The third assumption is that there are no gene-environment interactions or gene-environment passive correlations. Based on the gene-environment interaction, different sources of bias are expected. If AxC is present, then 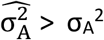 is expected. If AxE is present instead, 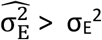If passive *r*_*G,E*_ is present, then 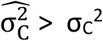 is expected. An additional set of assumptions introduced when estimating parameters via SEM is that means and variances are equal across zygosity group, twin order (i.e., 1 and 2), and sex. Details on the latter set of assumptions are given below. Complex sources of upward or downward biases in CTD-informed models (e.g., heterogeneity) are discussed elsewhere ^78^.

#### Baseline model

We first fit multigroup SEM models to create a baseline against which to compare the fit of univariate and multivariate models and test for the assumptions of the equality of mean and variances. The models freely estimated all the observed variance and covariances and included the age of the twins as a covariate. For the univariate model, equality of means and variances was tested by sequentially constraining parameters and comparing the Akaike Information Criterion (AIC) and Bayesian Information Criterion (BIC) of the model to the baseline model, where 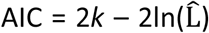and 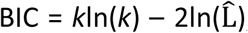 with *k* being the number of parameters estimated in the model and 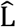the maximised value of the likelihood function. Models with smaller AIC and BIC than the baseline model were deemed a good fit. Additional comparisons are provided by the likelihood-ratio test (LRT), using the lavaan::lavTestLRT() function from the lavaan R package ^79^. All models were specified following lavaan notation and fitted with the lavaan::sem() function.

#### Univariate variance decomposition

The SEM specification was informed by the CTD, following the pattern of twin pairs correlations extracted from the baseline model and baseline model comparison results. Twin pairs correlations were extracted using the most parsimonious constrained baseline model using the lavaan::standardizedSoultion() function. Precisely, we fit a five-group ADE sem model, where the five groups were formed by either full or incomplete MZ female, MZ male, DZ female, DZ male, and DZ opposite-sex pairs. Means for women and men were estimated freely across sex, but not across zygosities or twin order. We fit the model via the direct symmetric approach by directly estimating the variances, as it can derive asymptotically unbiased parameter estimates and is, therefore, less prone to type I errors ^80^. We then decomposed the variance-covariance matrix **T** of twin pairs into the **T**= **A** + **D** + **E** variance covariances, which was predicted as follows:

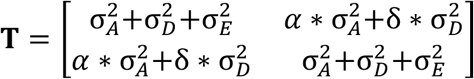

where *α* is the expected additive genetic relationship, and *δ* is the expected dominant genetic relationship between pairs (i.e., α = 1 or .5 δ =1 or .25, for MZ and DZ, respectively). Note that for simplicity, here we exclude the contribution of age to **T**, which was instead included in the model. To test for the significance of the variance components A and D, we additionally fit two models where D and AD variances were constrained to 0. Significance was inferred by model comparison, as above. We fit the model to the raw sum score of the BMRQ using the lavaan::sem() function. Assuming data within pairs were missing at random, we used the recommended estimator for twin data analysis, the full information maximum likelihood (FIML; argument estimator = “ML”). We used the following estimator for the narrow-sense heritability:

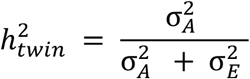

Here we note the detail that σ_A_^2^ + σ_E_^2^ ≠ σ_P_^2^, as σ_P_^2^ = σ_A_^2^ + σ_E_^2^ + B^2*^σ_Age_^2^. We also note that, since the E component includes residual deviation, σ_E_^2^ = inter-σ_E_^2^ + intra-σ_E_^2^, where inter-σ_E_^2^ is the inter-individual variance, and intra-σ_E_^2^ is the intra-individual variance ^77^. Comparisons with standard OpenMX protocols are given in Supplementary Note 7 (note that the small differences in test statistics did not lead to different conclusions). A graphical representation of the full univariate multigroup model can be found in Supplementary Fig. 5.

#### Sequential multivariate model

The sequential multivariate modelling of SMDT, BAS-RR, and BMRQ twin data was inspired by the classical multivariate Cholesky decomposition of additive genetic (A) and environmental (E) matrices ^81^. Following the CTD, we specified a multivariate model to estimate variance components and between-components between-trait path coefficients, *λ*_*A*_ and *λ*_*E*_, based on the between-trait between-twin (also referred to as cross-trait cross-twin) covariances. However, since variance components are directly estimated, it is important to note that the sequential multivariate model is not exactly a Cholesky decomposition. In fact, the predicted A and E variance-covariance matrices are not obtained as **A** = **XX**^T^ or **E** = **ZZ**^T^, as in a Cholesky decomposition, where **X** and **Z** are the lower triangular matrices with the path coefficients for the additive genetic and environmental components. Instead, the 6x6 variance-covariance matrix **S** was decomposed into symmetric matrices as **S** = **A** + **E**. As for the univariate case, the 6x6 symmetric matrices **A** and **E** include the predictions for the phenotypic variances and the twin pair phenotypic covariances. For comparison, we provide parameter estimates derived from the standardised solution, which is equivalent to a Cholesky decomposition, in Supplementary Fig. 6. Additionally, the **S** matrix also included the predictions for the within-twin and the between-twin between-trait covariances. One important consequence of our model specification is that we do not impose an implicit lower bound of zero on the variance components, which can cause bias when comparing different models. The sequence of variables was purely chosen to regress out A_1_ and A_2,_ respectively, implied from SMDT and BAS-RR observed scores, from the BMRQ. To estimate an adjusted heritability (here, for simplicity, adj-*h*^2^_twin_), we calculated the proportion of variance of the BMRQ covarying with the component A over the total BMRQ variance (minus the variance in BMRQ covarying with age):

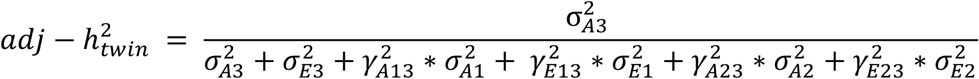

Where the numerical subscripts simply indicate the order of phenotype in the model (e.g., 3 is the BMRQ). To calculate the amount of additive genetic variance unique and associated with BMRQ beyond SMDT and BAS-RR (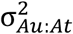, u=unique, t=total) we computed the proportion of genetic variance over the total BMRQ additive genetic variance as follows:

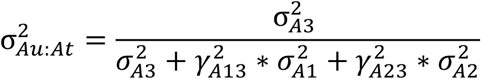

A graphical representation of the full multivariate model can be found in Supplementary Fig. 7. Similar to what was reported above, we fit the models using the lavaan::sem() function (estimator “ML”).

### Distinct factor solution

To estimate the genetic and environmental correlations between facets of music reward, we applied a correlated factor model via direct symmetric approach ^80^ (referred to as distinct factor solution). The direct symmetric approach is conceptually similar to a correlated factor solution. In the correlated factor solution, the multivariate phenotypic variance-covariance matrix **M** is obtained as **M** = **A**+**E** (in the simplest case of an AE model), with **A** =**XR**_A_**X**^T^ and **E** = **ZR**_E_**Z**^T^, where **X** and **Z** are the diagonal matrix of the standard deviation σ_A_ and σ_E_ and **R**_A_ is the genetic correlation matrix. Within a direct symmetric approach, instead, a different parametrisation is specified to directly estimate the **M** 10x10 symmetric matrix as **M** = **A** + **E**:

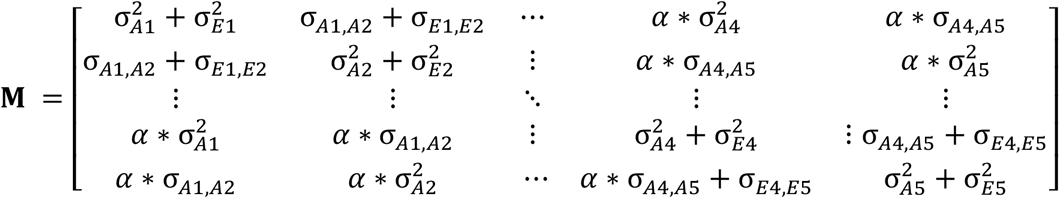

Where the **M**_1:5,1:5_ and **M**_5:10,5:10_ elements include the within-twin variance and between-traits covariances and are constrained to equal across zygosities, and the M_5:10,1:5_ and M_1:5,5:10_ elements include the between-twin additive genetic within- and between-trait covariances and the expected additive genetic relationship *α*, which is fixed to either 1 or .5 in MZ and DZ groups, respectively. While this approach may return out-of-bound values, the absence of boundaries has been shown to yield asymptotically unbiased parameter estimates and correct type I and type II error rates ^80^. A graphical representation of the full multivariate model can be found in Supplementary Fig. 8. Model syntax was written following lavaan specifications. Model fitting was done via the lavaan: sem() function (estimator “ML”). In sum, the distinct factor solution provides a multivariate model for the decomposition of phenotypic variances and covariances in genetic and environmental components. Comparison of this model with more parsimonious independent pathway models allows us to test for the presence of a common genetic (or environmental) component shared across facets.

#### Common-genetic factor solution

The hybrid independent pathway model (referred to as common-genetic factor solution) is a multivariate approach similar to the correlated factor solution, except with an additional restriction on the genetic covariances between traits (σ_A,A;_ hence hybrid or genetic, as environmental covariances are modelled in a distinct factor solution fashion). Consider a 5x5 phenotypic variance covariance matrix **P**. Under a hIPM AE model, **P** can be written as **P** = **A**_**c**_ **+ A**_u_ + **E**, where **A**_**c**_ = **X**_**c**_**X**_**c**_^T^, with **X**_c_ being a 5x1 vector of the additive genetic path coefficients of a common additive genetic factor (A_C_) loading across all phenotypes, and **A**_u_ is a 5X5 diagonal matrix including the residual unique genetic variance for each phenotype, σ_Au_^2^. The full additive genetic variance-covariance matrix can be then as follows:

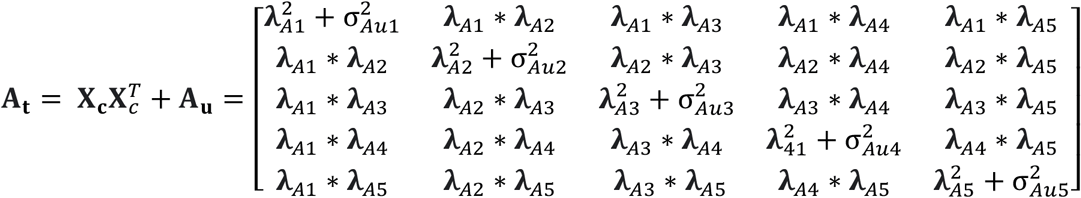

The 5X5 residual environmental covariance **E** simply contains the unconstrained residual environmental variances and covariances σ_E_^2^ and σ_E,E_. The 10X10 between-facet between-twin matrix **M** can then be written as follows:

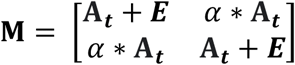

Where *α* is the expected additive genetic relationship between twins and is fixed to either 1 or .5 across MZ and DZ groups, respectively. A graphical representation of the full multivariate model can be found in Supplementary Fig. 9. Model syntax was written in lavaan. Model fitting was done via the lavaan:sem() function. Model comparison between distinct and common-genetic factor solutions was carried out via the laavan:: lavTestLRT() function. Here, we additionally note that the common-genetic factor solution is a less parsimonious version of the more commonly used independent pathway model and, therefore, provides a less restrictive and more specific test for a genetic common factor when compared to the distinct factor solution.

#### Structural equation modeling assumptions

SEM-based estimates obtained from the full information maximum likelihood (FIML) estimator are unbiased under the assumption that observations follow a multivariate normal distribution ^41^. Violation of the assumption of multivariate normality has been found to have little impact on parameter estimates but can have severe consequences for both the χ^2^ test statistics and the standard error of the estimates for the parameters. An alternative estimator that is less sensitive or robust to violation of multivariate normality is the maximum likelihood with robust standard error and scaled test statistics (MLR). Although this estimator assumes missingness to be completely at random, it has been shown to provide quite reliable estimates of data missing at random ^82^. Relevant comparisons between the two estimators are given in Supplementary Note 1.

## Data availability

The datasets generated during the current study cannot be made public as registry data were used. However, researchers are able to apply online at the Swedish Twin Registry to access the twin data used in this study (see https://ki.se/en/research/swedish-twin-registry-for-researchers).

## Code availability

All scripts and code used to analyse the data can be found at: https://github.com/giacomobignardi/h2_BMRQ.

## Acknowledgements

We thank the Swedish twins for their participation. The Swedish Twin Registry is managed by Karolinska Institutet and receives funding through the Swedish Research Council under the grant no 2017-00641. We further thank Anirudh D. Patel for his critical feedback and Cristina Gonzalez-Liencres for her input on earlier versions of the figures and figures’ captions. We also would like to thank the participants of the 2023 Nijmegen *Musicality Genomics Consortium* meeting (https://www.mcg.uva.nl/musicgens/program.html) for insightful discussions that have led to an improved version of the current work. G.B. was supported by the German Federal Ministry of Education and Research (BMBF); G.B. and S.E.F. were supported by the Max Planck Society.

## Contributions

G.B. conceived the study, analysed and visualised the data; G.B. and M.M. drafted the manuscript; F.U., S.E.F., and M.M. supervised the research; L.W.W. validated the work; S.E.F, M.M., R.J.Z., L.W.W., and F.U. conceptually validated the work; all authors revised and reviewed the last version of this manuscript.

